# Mate sharing in male stump-tailed macaques as a possible case of coalition-like behavior to modify the group-wise fitness distribution

**DOI:** 10.1101/2020.01.30.927772

**Authors:** Aru Toyoda, Tamaki Maruhashi, Suchinda Malaivijitnond, Hiroki Koda, Yasuo Ihara

**Author notes:** **Correspondence to**: Aru Toyoda, Yasuo Ihara.

## Abstract

In multi-male multi-female groups of animals, male reproductive success is often skewed toward top-ranking males. Coalition formation by middle- to low-rankers can be seen as a collaborative effort to modify the distribution of reproductive success within the group, so that they gain more than they would do otherwise. It has been pointed out, on the other hand, that a coalition of top-ranking males could hardly be profitable in the sense that they would gain little additional benefit from making collaborative effort. Here we report our novel observation of facultative sharing of mating opportunities among males in a wild population of stump-tailed macaques (*Macaca arctoides*) as a possible case of coalition-like behavior in which dominant males jointly guard females from mating with subordinate males and actively share mating opportunities within the allies. First, we report our novel observation of facultative sharing of mating opportunities in male stump-tailed macaques, where two or more males remain in close proximity to and copulate with a female in turn without contesting or sneaking. Second, considering the kind of coalition formation in which dominant males collaboratively exclude subordinates from mating competition and thereby strengthen the reproductive skew that already exists, we specify, by means of mathematical modeling, the condition for this kind of coalition formation to be selectively favored. Finally, we derive predictions about the occurrence of the coalition-like behavior depending on ecological and demographic factors, and test them empirically using data from the five groups of stump-tailed macaques in our study population.

---

Male reproductive success in a multi-male multi-female group is often skewed in favor of high-ranking males (Kutsukake & Nunn, 2006). As a countermeasure, low-ranking males may collaborate to modify the distribution of reproductive success within group so that they gain more than they would do otherwise. Male collaboration that results in modification of within-group resource allocation has been theoretically investigated in the context of coalition or alliance formation (Pandit & van Schaik, 2003; van Schaik et al., 2004, 2006). Assuming that a male’s share of mating opportunity decreases monotonically with descending rank position, Pandit & van Schaik (2003) developed a mathematical model of leveling coalition, in which a skew in the access to females in favor of dominant males is mitigated by coalition formation of middle-to low-ranking males. While animal coalition is typically manifested as a coordinated aggression by multiple individuals on one or more targets (Bissonnette et al., 2009; Smith et al., 2010) or an individual’s intervention in an ongoing conflict between two parties to support one side (Widdig et al., 2000), the Pandit-van Schaik model is potentially applicable to any case where male-male collaboration serves to flatten the group-wise fitness distribution, even in the absence of overt expression of coordinated aggression.

In Pandit & van Schaik’s (2003) terminology, a coalition of males ranking below the target (i.e., revolutionary or all-up coalition; (Chapais, 1995; van Schaik et al., 2006) is profitable in the sense that it confers, if successful, a fitness benefit to all coalition members. Hence, whenever the coalition is also feasible, that is, if the subordinates are able to jointly beat the dominant target, this type of coalition is predicted to occur. In contrast, a coalition of top-ranking males against a lower-ranker (i.e., conservative or all-down coalition; (Chapais, 1995; van Schaik et al., 2006) would not be profitable because each of the top-rankers can beat the target even without help. A simplest implication from the Pandit-van Schaik model is, therefore, that male-male collaboration to modify the group-wise distribution of reproductive success is more likely to occur among middle-to low-ranking males than among top-rankers (Pandit & van Schaik, 2003; van Schaik et al., 2004, 2006). The expectation is at least partially supported in some species of primates (Bercovitch, 1988; Noë & Sluijter, 1995), but not in others (Young et al., 2014).

In this article, we report a novel observation of facultative sharing of mating opportunities among males in a wild population of stump-tailed macaques (*Macaca arctoides*). We regard the observed mate sharing as facultative because there the males copulate with a female in turn without contesting or sneaking. As an illustration, Fig. 1 shows two male stump-tailed macaques sharing a female without exhibiting any sign of antagonism, where the male on the right is copulating with the female while the male on the left is waiting for his turn. In some, but not all, groups in our study population of stump-tailed macaques, the alpha (i.e., the highest-ranking) male forms a coalition-like unit with one or more other males in the same group. The allied males seem to jointly guard a female from mating with other males, while within the coalition-like unit they share the mating opportunities.

**Figure 1.**
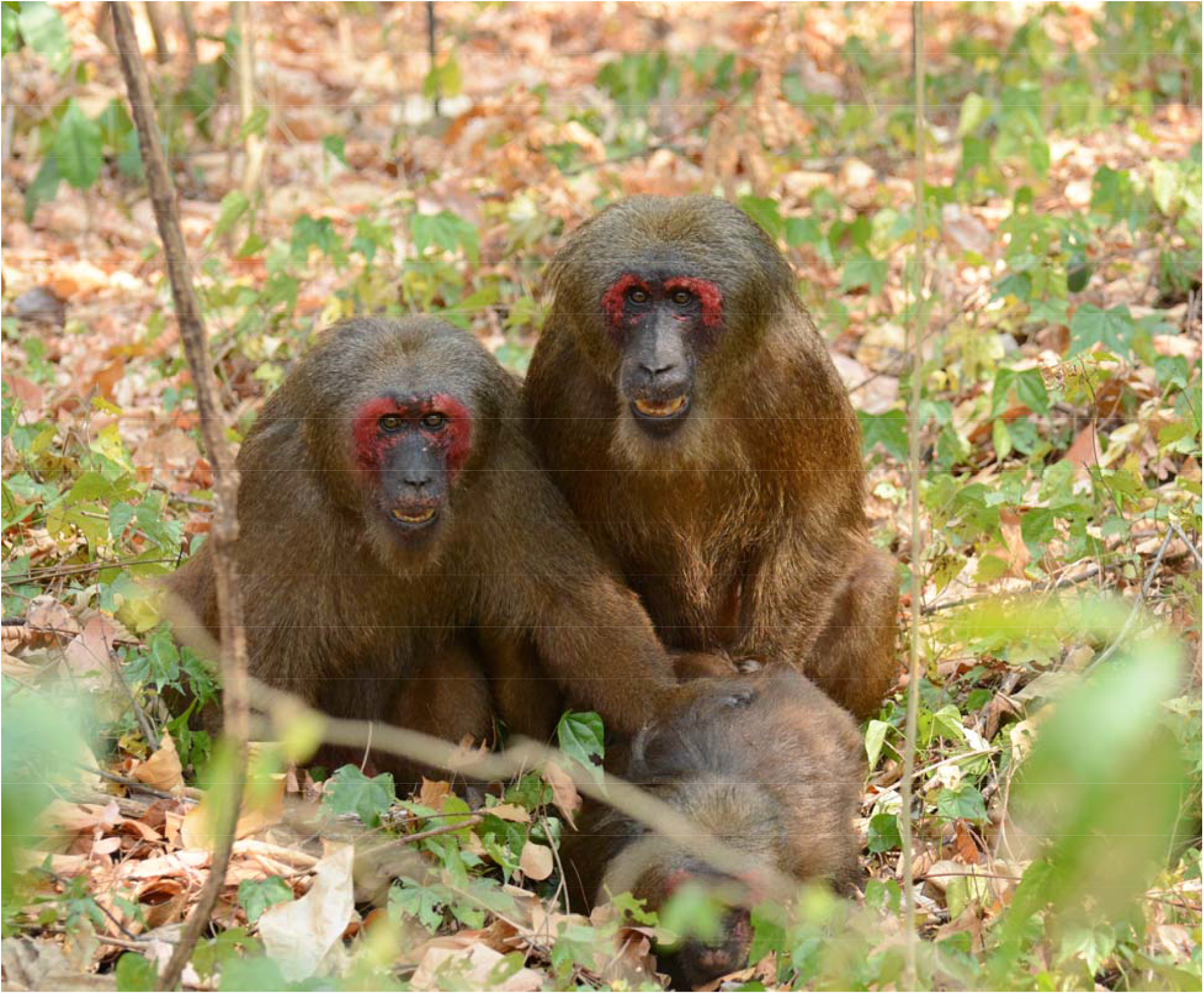
Stamp-tailed macaque males (FTH-M01 and FTH-M02) sharing mating opportunities with a female (FHT-F11) without showing any sign of antagonism.

Strictly speaking, the observed collaborative unit of males is not to be called a “coalition” as it lacks an overt expression of coordinated aggression; nonetheless, we choose to refer to it as “coalition like” because it is a form of male-male collaboration that can potentially modify the group-wise distribution of male reproductive success, as the genuine coalition does in the Pandit-van Schaik model. In particular, suppose that the number of sexually receptive females in a group is larger than that would allow the alpha male to monopolize reproduction (i.e., complete contest competition), but smaller than that would lead to complete scramble competition. In such cases, top-ranking males may be able to exclude subordinates more efficiently from mating competition by guarding females collaboratively than by doing so individually. This raises the possibility that dominant males may gain a fitness benefit by having collaborative partners despite the cost of sharing mating opportunities with them.

Since mate sharing in stump-tailed macaques occurs around the alpha male, it is similar to all-down coalition. Within the framework of Pandit and van Schaik’s (2003) leveling coalition, however, all-down coalition is not predicted to occur. Instead, the mate sharing by dominant males may be better represented as a novel kind of coalition formation that deprives subordinate males of mating opportunities, whereby “steepens” the reproductive skew. For the purpose of investigating the coalition-like behavior in stump-tailed macaques, we extend the Pandit-van Schaik model by allowing all-down coalition to enhance the efficiency with which top-ranking males keep subordinates from mating, as a result of which the reproductive skew is strengthened. To keep the model simple and tractable, we do not explicitly specify the underlying mechanism by which subordinates are excluded. A recent study by Pandit, Pradhan, & van Schaik (2020) took a different approach to a similar question. They extended the original Pandit-van Schaik model by incorporating a specific mechanism to realize more skewed resource allocation, namely, higher-rankers’ usurping of resources owned by lower-rankers. However, since Pandit et al.’s (2020) model is designed to explore the origins of class formation in human societies, and non-human primates are unlikely to meet the presumption that individuals possess exploitable or tradable resources, it is not applicable to the current context.

In what follows, we describe our observation of facultative mate sharing among male stump-tailed macaques, develop a mathematical model to explore the underlying mechanism of the behavior, and empirically test predictions derived from the model.

## METHODS

### Study site and animals

A wild population of stump-tailed macaques inhabiting the Khao Krapuk Khao Taomor non-hunting area in the Phetchaburi Province of central Thailand (99°44’ E, 12°48’ N, encompassing an area of 3.5–4 km^2^) was observed. This site consists primarily of secondary forests, including stands of bamboo and agricultural areas. The macaques also visited areas immediately adjacent to this site (including a nearby temple, cassava and pineapple plantations, and human settlements) on a daily basis. The macaques were occasionally fed by humans, both locals and tourists, on the temple grounds or along the roadside. This population was first documented in June of 1984, at which time there were only 22 individuals. Since then, it has grown to a large population, including at least 391 individuals (97 adult males, 124 adult females, 114 infants (≤ 2 years), and 56 subadult unidentified subjects, who were divided into five groups, namely, Ting, Nadam, Third, Fourth, and Wngklm groups (Table S1 in Supplementary Materials) by 2017. The Wngklm group separated from the Third group in November−December 2015. All adults (completely mature monkeys), most subadults (sexually mature but not completely developed), and some juveniles (sexually immature, around 3 years of age) were identified based on facial characteristics. This population is geographically isolated from the other populations, and no new immigrant males from other sites were detected during the study period (Toyoda et al., 2017; Toyoda & Malaivijitnond, 2018).

Stump-tailed macaques are reported in general as non-seasonal breeders (some local populations/captive groups show seasonality, but that is not the case for the populations in Thailand). No clear sign of estrus is observed in females, and thus it is not possible, at least for human observers, to detect ovulation. Concealed ovulation is generally considered to affect the distribution of male reproductive success by hindering the alpha male from monopolizing fertile females. It should be noted, however, that there is an ongoing debate about whether or not males can infer reproductive status of females. One paper has suggested that males can detect female ovulation by vagina testing behavior (Cerda-Molina et al., 2006) although this is not congruent with our data suggesting that more than half of copulations occurred in non-fertile period (Toyoda, unpublished data).

### Daily observations

AT performed 21-month field observations for the five groups, between September 25, 2015 to June 15, 2017. In total, the animals were observed for 289 days (970.7 hours). The monkeys were followed daily between 09:00 and 17:00 h; the group that was first encountered each day was followed for as long as possible. When the target group could not be followed further (e.g., when the monkeys travelled along cliffs), the observation of the target group was terminated, and another group was sought out and followed.

All copulations during our observation were recorded using video cameras (JVC GZ-RX500 and Sony HDR-PJ675) and their descriptions were noted.

### Data on copulation

Following a previous report (Estep et al., 1984), *single copulation* was defined as a sequence of copulating behavior consisting of mount, insertion, and separation, irrespective of the presence or absence of ejaculation. The starting and ending times of each single copulation were recorded, and the inter-copulation intervals (ICIs) of all adjacent events of single copulations recorded within a day were calculated. A series of single copulations by a female at an interval of less than 30 min was considered collectively as a *serial copulation bout* if it included four or more single copulations. We adopted this definition for its improved objectivity as compared with the relatively ambiguous definition in previous studies (Brereton, 1994; Estep et al., 1984). Each of all single copulations not included in any serial copulation bouts was considered a *non-serial copulation bout*.

### Mathematical model

To understand the logic behind the coalition-like behavior in male stump-tailed macaques, we designed a simple mathematical model based on the framework developed by Pandit & van Schaik (2003). Our motivation for the mathematical modeling was two-fold. First, it is intuitively conceivable that concealed ovulation, as in stump-tailed macaques, hinders the alpha male from guarding all fertilizations, and thus necessitates collaboration of two or more dominant males for reproductive monopoly. However, it is unclear as to whether and under what circumstances the alpha male tolerates one or more allies copulating with females. Second, we observed within-species variation in the occurrence of the coalition-like behavior: it occurs in the Ting, Nadam, and Fourth groups, but not in the Third and Wngklm groups (see Results section). If our logic is sound, the model should also explain this pattern of within-species variation.

We consider a group of *N* males and a constant number of females. In the case of a linear order of dominance among the males, the relative access of the *i*th male to females, *x*_*i*_, in the absence of male-male coalition is described by the priority-of-access model (Altmann, 1962), namely,

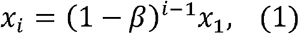

where *β* represents the degree to which dominant males can monopolize mating opportunities, called the despotic parameter (0 < *β* < 1).

Ecological and demographic factors have been suggested to affect *β* (van Schaik et al., 2006). Of these, cryptic ovulation in females probably reduces *β* as it prevents dominant males from guarding each female intensively only during her fertile period. Thus, compared to species in which ovulation is advertised, species with cryptic ovulation are expected to have smaller *β*. Other factors, such as the number of females in the group, the relative strengths of dominant males, and the female preference for or against dominant males, are also likely to affect *β*. As the latter factors may vary within a species, we expect that different groups of stump-tailed macaques are characterized by different *β* values.

There may exist circumstances such that it is beneficial for top-ranking males to form a coalition to guard potentially fertile females in a collaborative manner and share copulation opportunities among the allies. The effect of this behavior on the distribution of male reproductive success can be represented by the following equation:

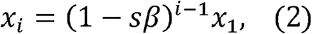

where we assume 1 < *s* < 1/*β*. Parameter *s* reflects the effect of male-male coalition to “steepen,” instead of “flatten,” the male reproductive skew, where larger *s* indicates higher reproductive monopolization by dominant males. It should be emphasized that our parameter *s* differs from *α*, the similar parameter in the Pandit-van Schaik model, the latter of which captures the effect of coalition among subordinate males to level the reproductive skew (i.e., 0 ≤ *α* ≤ 1).

To evaluate the profitability of a male-male coalition, the cost and benefit of collaboration has to be defined. We considered two components of a particular male’s fitness: the ratio of the mating opportunities gained by that male to all the existing mating opportunities, which approximates the male’s share of paternity, and the cost associated with the additional effort of collaborating with others. As for the first component, *y*_*i*_ denotes the proportion of mating opportunities obtained by the *i*th male among all matings; that is,

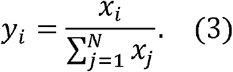

From (1) and (2), in the absence of male-male coalition, we obtain

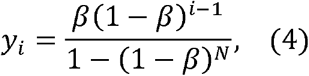

while in the presence of a coalition,

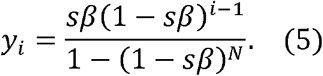

Regarding the second component, we assume that the first components of males forming a coalition are multiplied by 1 − *c*, where *c* represents the cost of collaboration (0 < *c* < 1), while those of non-coalition males are multiplied by 1. Therefore, for the *i*th male, joining a coalition is profitable if and only if

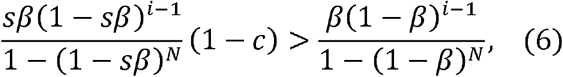

or equivalently,

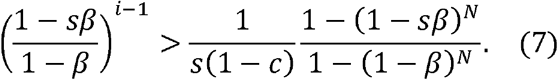

As 1 − *sβ* < 1 − *β*, the left-hand side of (7) decreases with increasing *i*; thus, whenever (7) holds for the *m*th male (*m* ≥ 2), it also holds for the first to *m* − 1th males.

### Estimation of parameters

We estimated the despotic parameter, *β* in (1) or *sβ* in (2), using nonlinear regression analysis on the observed distribution of the number of copulations, as a proxy to the reproductive success, among males in each of the five groups. For the purpose of better approximating male reproductive success, we considered only those single copulations for which ejaculation was confirmed. Males were sorted according to the number of copulations by the descending order, and a rank order, *i*, was assigned to each of them. We used the rank among males in the number of copulations as a substitute for the dominance rank in the Pandit-van Schaik model, assuming that coalition formation does not change the dominance ranks of the coalition members of the target. The *i*th male’s number of copulations, *x*_*i*_, was regressed on the rank order, *i*, to estimate *β* in (1) or *sβ* in (2) using the nonlinear regression function of Python (curve_fit method in SciPy optimize module). In addition, *R*^2^ values were reported for showing the goodness-of-fitting.

### Ethics approval

All data acquisitions and procedures during the fieldwork were approved by the National Research Council of Thailand (NRCT, Permission No. 0002/6910) and the Department of National Parks, Wildlife and Plant Conservation of Thailand (DNPT). We also complied with the guidelines for field studies of the Primate Research Institute, Kyoto University.

## RESULTS

### Description of the observed behavior

We counted 433 cases of single copulations (defined as a single mount-insertion-separation sequence, see Methods section) from the five subject groups. The median and range of the inter-copulation intervals (ICIs; *N*_*ICI*_ = 206) were 7 and 0-359 min, respectively, and 95% of the ICIs were 30 min or shorter (Fig. 2). Out of the 433 copulations, 213 (49.2 %) cases occurred in serial copulation bouts, while 220 (50.8%) occurred as non-serial copulation bouts. In total, 26 serial copulation bouts were recorded in the five groups (nine cases in the Ting group, four in Nadam, five in Third, six in Fourth, and two in Wngklm; for details see Table S2 and Fig. S1 in Supplementary Materials), among which the median and range of the per-bout number of single copulations resulting in ejaculation were 6 and 1-31 times, respectively.

**Figure 2.**
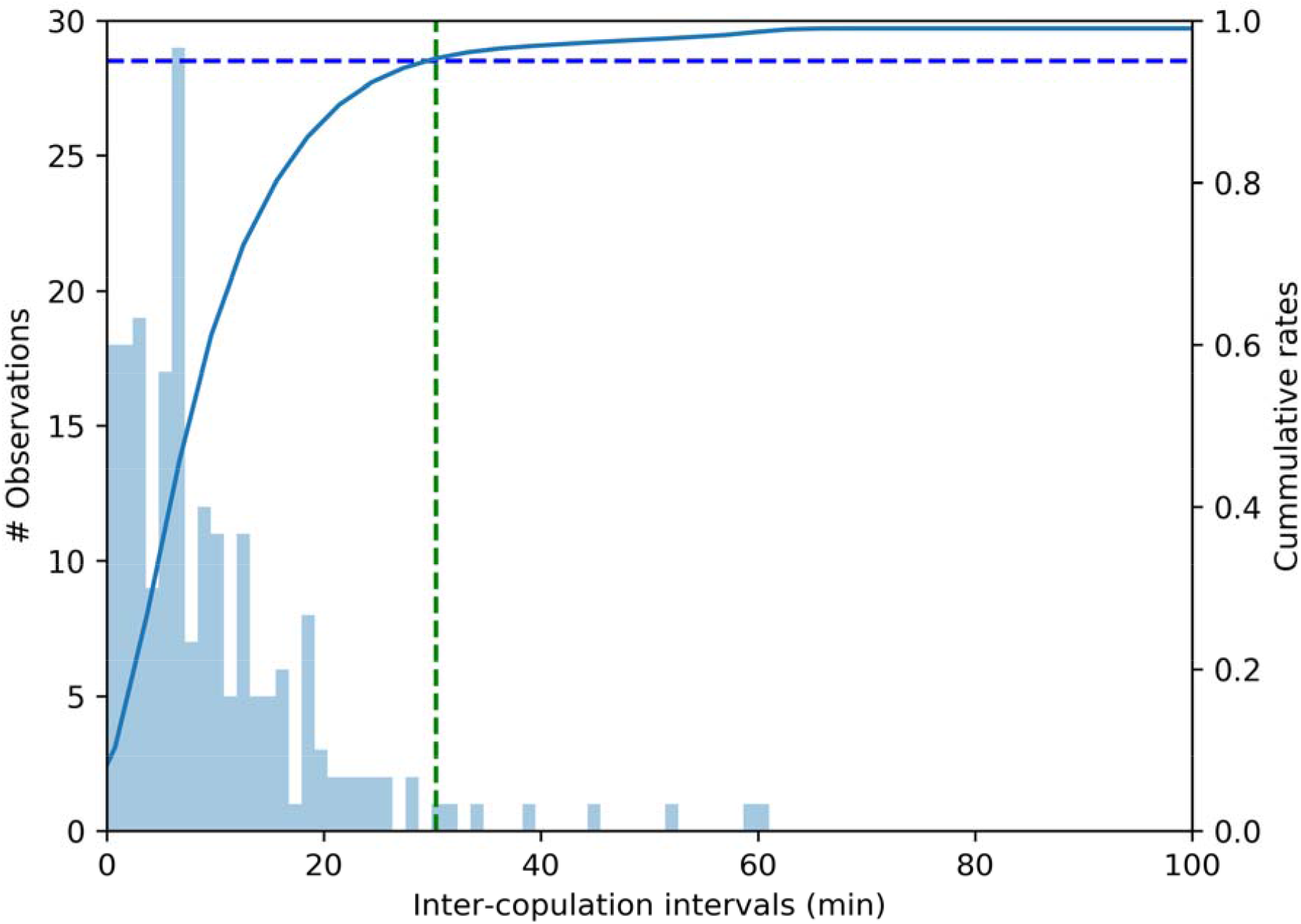
The distribution of the inter-copulation intervals (ICIs) between two consecutive single copulations that involved the same pair of male and female (*N*_lCl_ = 206). The green broken line represents the critical ICI value below which 95% of the observed ICIs were included. Note that only the cases where the same pair of male and female copulated more than once within a day are included.

In the Third and Wngklm groups, a single male was responsible for all of the five and two serial copulation bouts, respectively. These were the same males as those considered as the alpha males throughout the study period on the basis of behavioral observation (named TRD-M01 and WKM-M01 for the Third and Wngklm groups, respectively), i.e., in these groups the alpha males remained in the proximity of a female and copulated repeatedly. Pooling serial and non-serial copulation bouts, we recorded 80 single copulations by TRD-M01, of which 68 were associated with ejaculation, amounting to 81.8% of all copulations and 86.1% of all ejaculated copulations in the Third group. Similarly, we observed 26 single copulations including 22 ejaculatory copulations by WKM-M01, which were 59.1% of all copulations and 78.6% of all ejaculatory copulations in the Wngklm group. These indicate that the alpha males were able to obtain considerable portion of copulation opportunities on their own in the Third and Wngklm groups (Fig. 3, see “alpha-monopoly” type).

**Figure 3.**
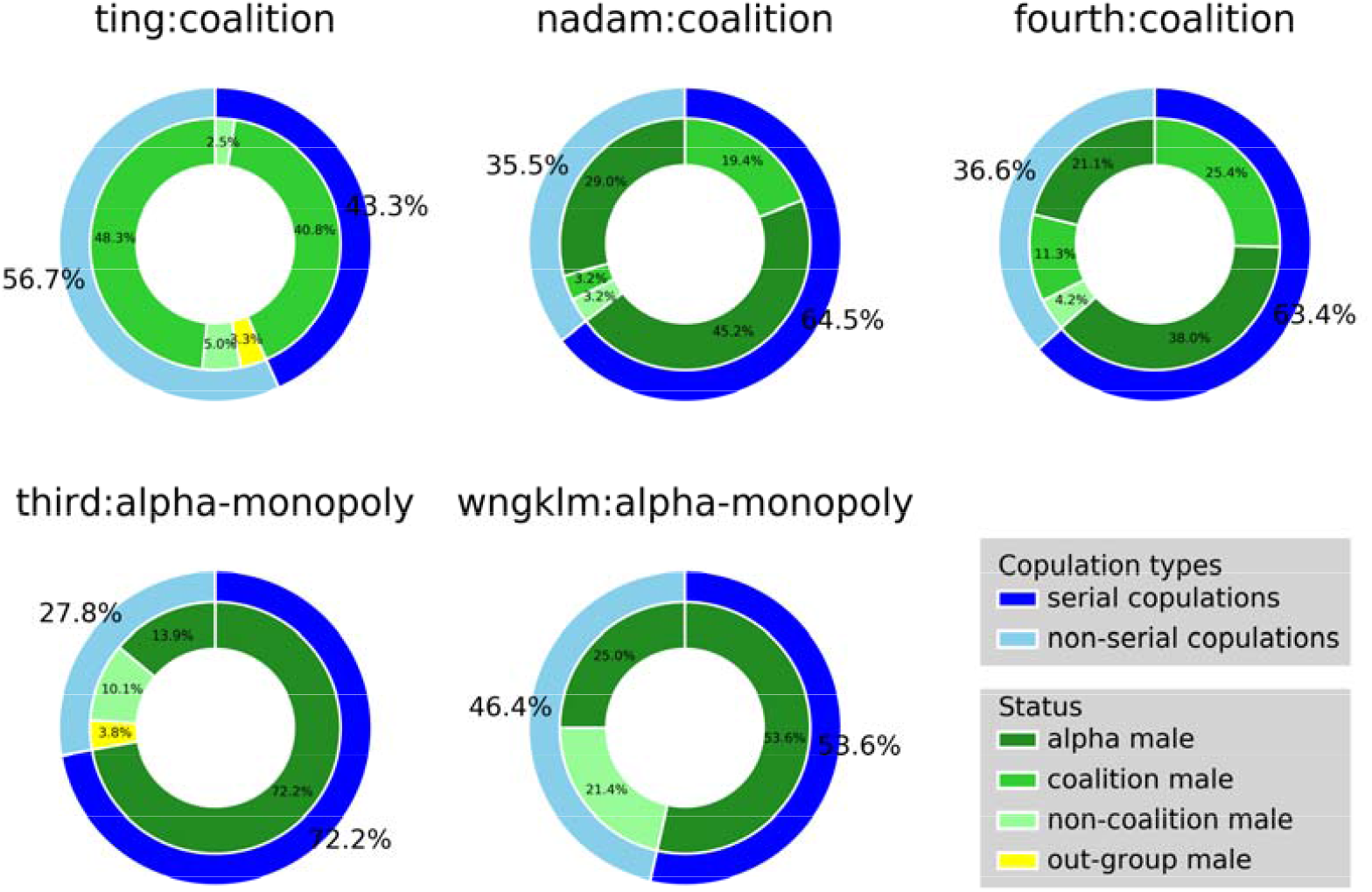
Proportions of single copulations with different characteristics to all single copulations in each of the five groups. Only are the single copulations associated with ejaculation included. The outer circle compares the proportions of single copulations that are part of serial copulation bouts and those that are non-serial copulation bouts. The inner circle shows the proportions of single copulations by the alpha males, non-alpha males forming a coalition unit, non-alpha males not forming a coalition unit, and males from outside of the group.

In contrast, we observed serial copulation bouts involving multiple males in the Ting, Nadam, and Fourth groups (Fig. 3, see “coalition” type). Interestingly, in each such multi-male serial copulation bout, males copulated with a female in turn without overt conflict; that is, while one male copulated with a female, the other male(s) maintained close proximity to the copulating pair, and only after one male performed several sequences of copulatory behaviors (i.e., mount, insertion, and separation), did another male take over the role as the copulator (see Fig. 1). In total, 14 cases of multi-male serial copulation bouts were observed, of which nine cases involved two males and five involved three males, and in no cases four or more males were involved (Table S2). The identities of the males sharing females were highly stable, particularly in the Nadam and Fourth groups, in which the numbers of males participated in at least one case of multi-male serial copulation bout were two and three, respectively, including NDM-M01 in Nadam and FTH-M01 in Fourth, who were considered as the alpha males throughout the study period. In the Ting group, where a clear dominance order was not established during the study period among several dominant males, there were nine males that participated in at least one case of multi-male serial copulation bout (Table S2). In addition, there were 108, 30, and 68 ejaculated copulations by those males that participated in at least one case of multi-male serial copulation bout in the Ting, Nadam, and Fourth groups, respectively, which amounted to 93.1%, 96.8%, and 95.8% of all ejaculated copulations excluding those done by outgroup males (Table S2, Fig. 3). This suggests that the copulation opportunities in the Ting, Nadam, and Fourth groups were obtained almost exclusively by those males who were members of the facultative mate sharing. From these findings, we hypothesize that those males sharing a female function as a coalition-like unit to jointly guard females from mating with other males and actively share the secured mating opportunities within the unit.

### Mathematical analysis

Our mathematical analysis specifies the condition for a coalition by the *i* highest-ranking males to be profitable (i.e., (7)). Since a coalition of top-rankers is always feasible, such a coalition is predicted to occur whenever (7) holds.

Regarding within-species variation in the occurrence of coalition formation, we investigate how ecological and demographic factors, which may vary among groups of stump-tailed macaques, affect the profitability condition (7). We derive from (7) the upper bound of coalition size, *m*^*^, for given *β, N, s*, and *c* as

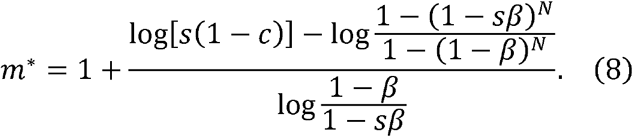

Equation (8) immediately shows that *s* (1 − *c* > 1 is necessary for any coalition to be profitable; otherwise, *m*^*^ < 1 always holds. It also shows that the right-hand side of (8) increases with *N* (Fig. 4a, 4b).

**Figure 4.**
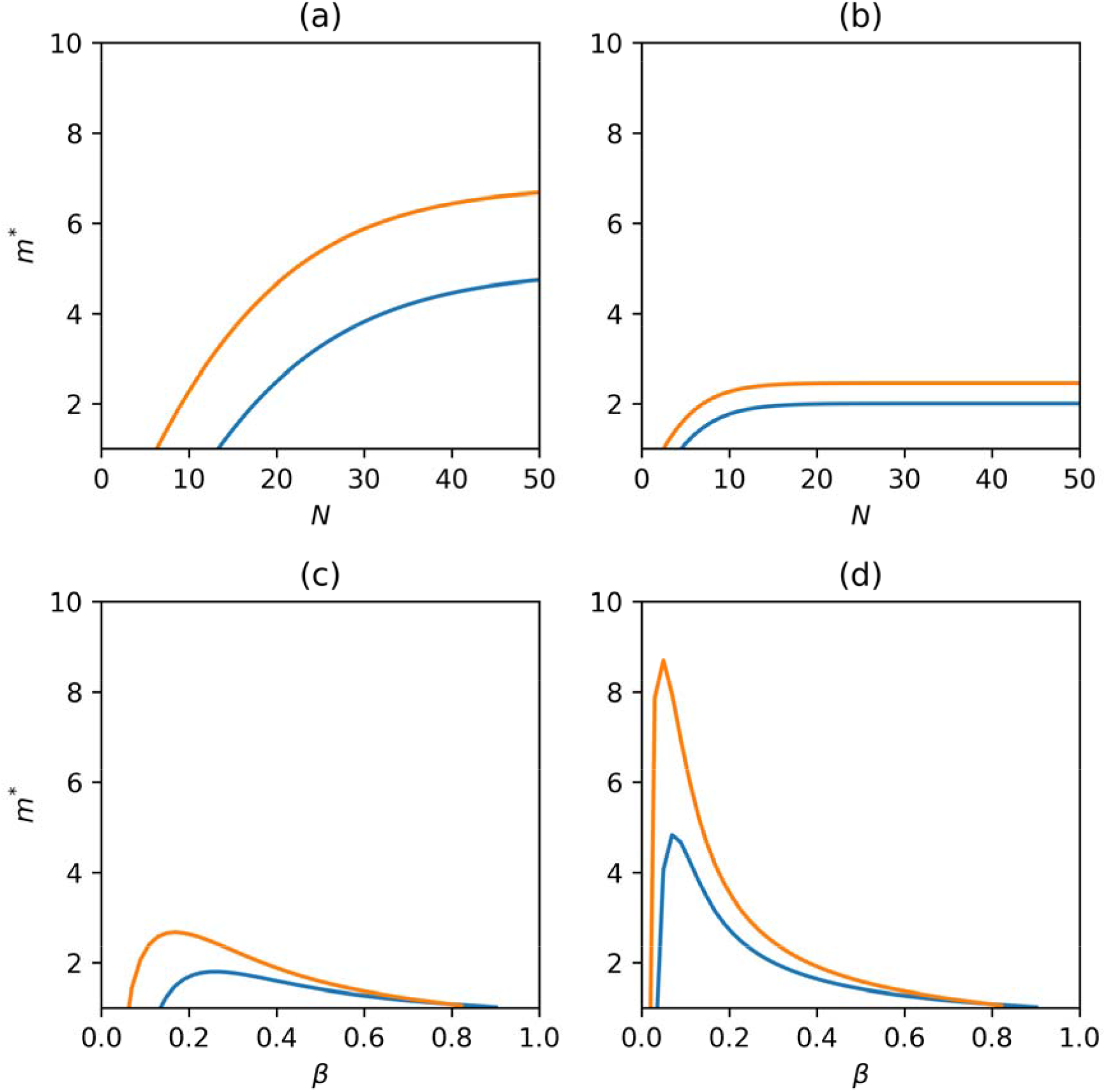
The dependence of the upper bound of coalition size, *m*^*^, on the number of males, *N*, and the despotic parameter, *β*, in the absence of coalition formation. The blue and orange curves represent *m*^*^ for *s* = 1.1 and *s* = 1.2, respectively. For all panels, *c* = 0.05. (a) *β =* 0.1, (b) *β =* 0.3, (c) *N* = 10, (d) *N* = 40.

For large *N*, the upper bound of the coalition size is obtained approximately as

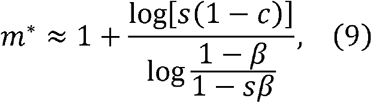

which decreases with increasing *β* whenever *s* (1 − *c*) > 1. On the other hand, when *β* is small, (8) is approximated by

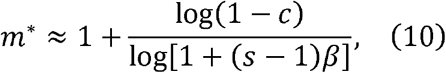

indicating that as *β* approaches zero, *m*^*^ diverges to minus infinity. In general, the dependenceof *m*^*^ on *β* is not monotonic (Fig. 4c, 4d). For coalition of at least two males (i.e., *m*^*^ > 2), (9) shows that *β* should be smaller than a threshold, specified by

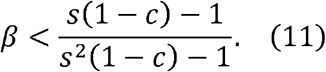

Hence, for any coalition to occur, *β* has to be relatively small, but not extremely small. Fig. 5 illustrates the combinations of *N* and *β* values, for which *m*^*^ > 2 (based on (8)).

**Figure 5.**
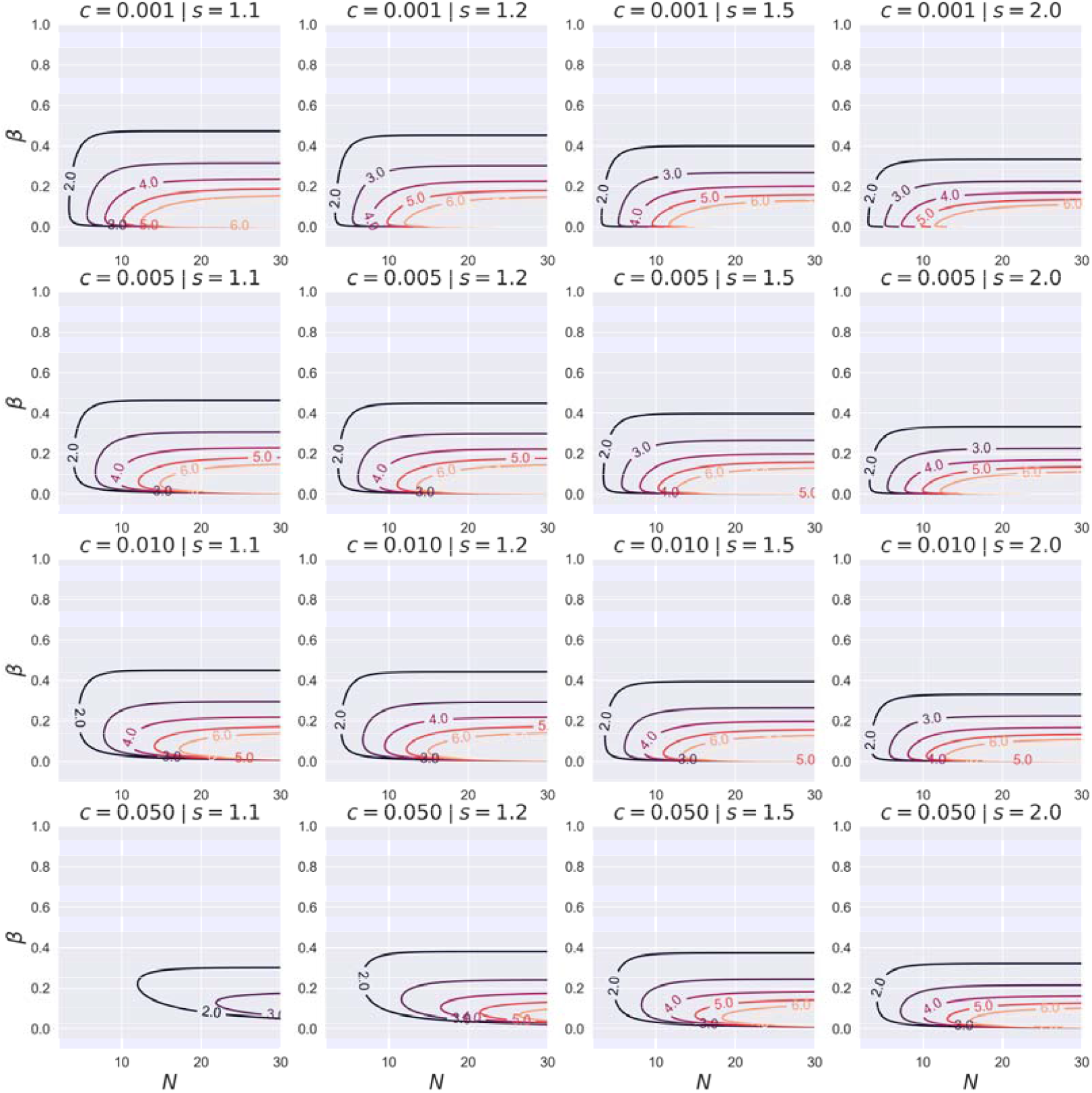
Contour plots of the upper bound of coalition size, *m*^*^, on the *Nβ* -plane for given values of *c* and *s*. Each contour represents (8) for the designated value of *m*^*^.

In summary, we have established the theoretical possibility of coalition formation by dominant males to steepen the reproductive skew that already exists, providing a formal foundation for our hypothesis of coalition-like behavior in male stump-tailed macaques. We have also specified the condition for this kind of coalition to occur, from which the following two predictions are derived: first, a large coalition is more likely to be observed in larger male groups (Fig. 5); and second, among sufficiently large male groups, a coalition is more likely to occur in a group where the extent of reproductive monopolization by dominant males is relatively small, unless it is extremely small (Fig. 5). If, as we have hypothesized, coalition-like behavior underlies the observed facultative mate sharing in stump-tailed macaques, patterns of within-species variation in the occurrence of the facultative mate sharing are expected to follow these predictions.

### Test of model predictions

We evaluated the above predictions for within-species variation on the basis of our observational data from the five groups of stump-tailed macaques. For each group, we estimated the despotic parameter, *β*′, that was supposed to be either unmodified (i.e., *β*′ = *β* for Third and Wngklm) or modified (i.e., *β*′ = *sβ* for Ting, Nadam, and Fourth) by male coalition-like behavior. Non-linear regression analysis returned the following estimates of *β*′: 0.30, 0.72, 0.97, 0.59, and 0.78 for Ting, Nadam, Third, Fourth, and Wngklm, respectively (Fig. 6).

**Figure 6.**
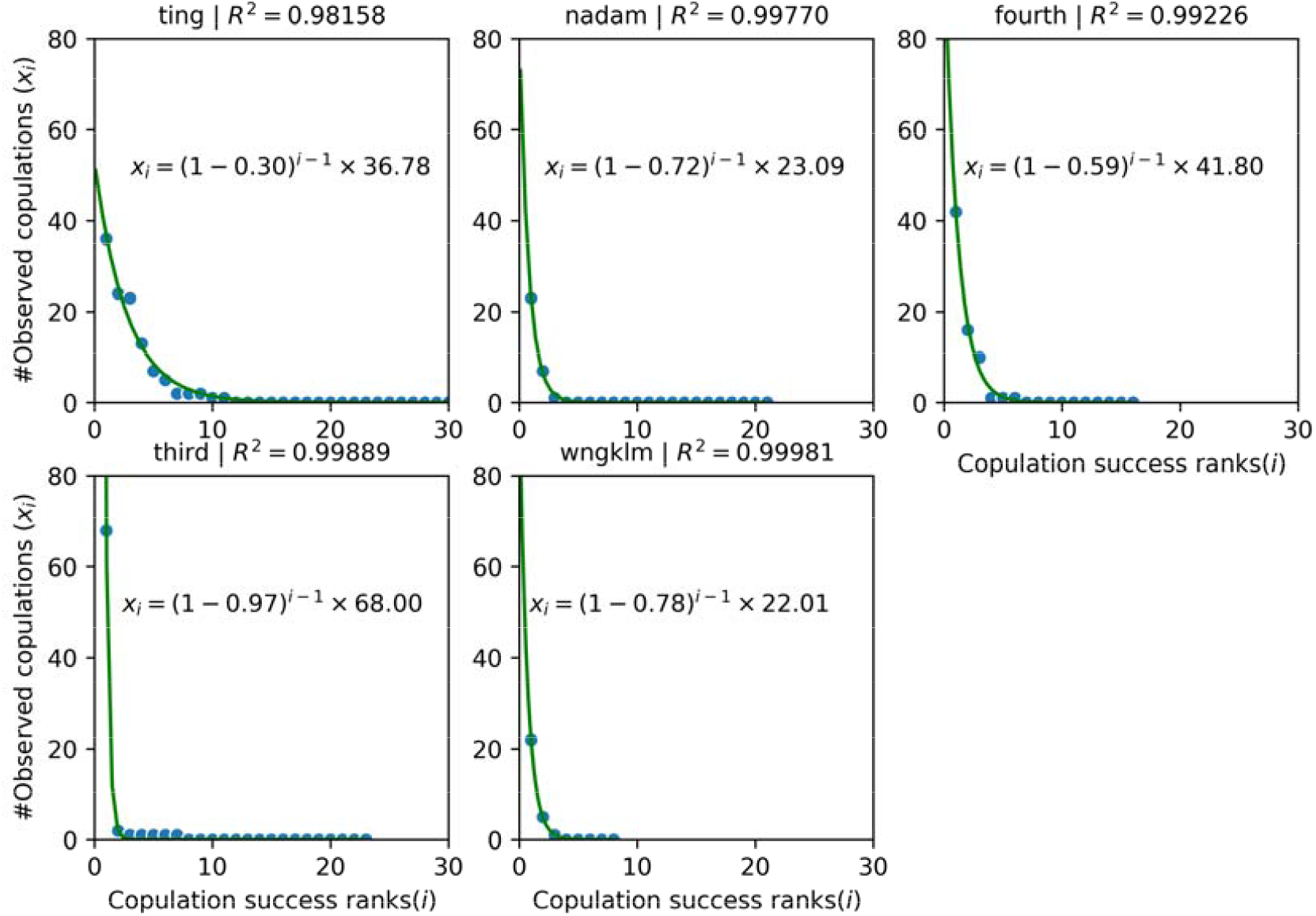
The observed distribution of the number of single copulations with ejaculation among males in each of the five groups. The curves represent the results of nonlinear regression analysis. *R*^2^ gives the coefficient of determination.

Figure 7 shows the estimates of *β*′ and the numbers of males, *N*, in the five groups. It is to be noted that the vertical axis represents *β* for Third and Wngklm and *sβ* for Ting, Nadam, and Fourth, so that the unmodified despotic parameter *β* in the latter three groups would be lower than these values. Consistent with the model predictions, the Ting group, in which there were nine males who participated at least one multi-male serial copulation bout (*m* = 9), had the combination of the largest *N* and the smallest *β*′ (and thus *β*) among the five groups. In addition, as predicted, Fourth (*m* = 3) and Nadam (*m* = 2), the two other groups in which multi-male serial copulation bouts were observed, had the second and third lowest values of *β*′ (and thus *β*), respectively. On the other hand, multi-male serial copulation bout is absent in the Third group with the second largest *N*, which might appear to contradict our predictions. We tentatively interpret this as a result of large *β*′ in this group; in other words, *β* may be too large to satisfy (11), although a quantitative evaluation of this claim has been challenging so far. In sum, we conclude that our model accounts well for the patterns of within-species variation in the coalition-like behavior in stump-tailed macaques.

**Figure 7.**
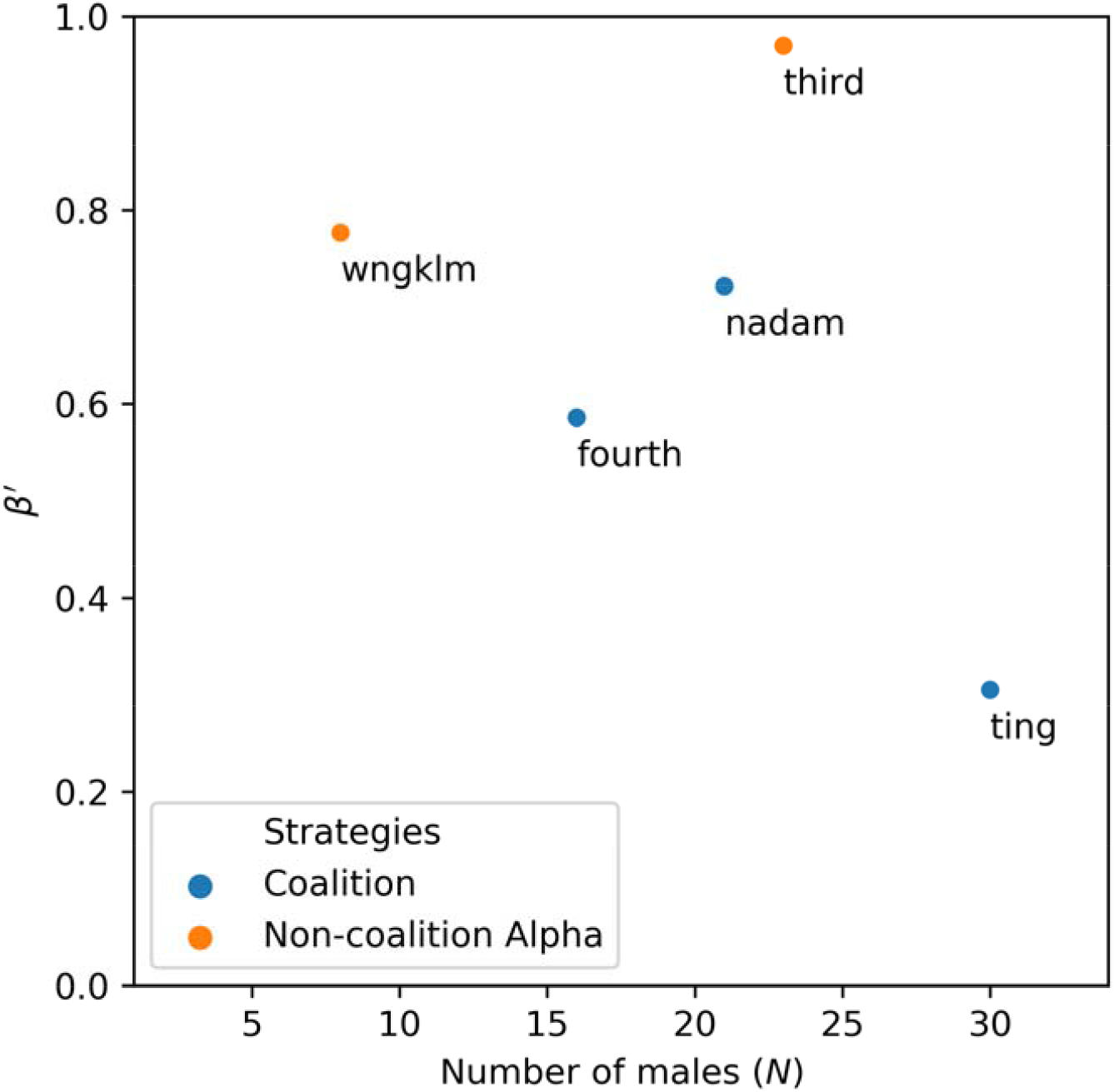
The estimates of *β*′ and the numbers of males, *N*, in the five groups.

## DISCUSSION

Our observations, together with mathematical analysis and empirical test of model predictions, suggest that the alpha male of stump-tailed macaques form a coalition-like unit with one or more males in the group to jointly secure mating opportunities and actively share them within the unit. To the best of our knowledge, this is the first mammalian observation of collaborative mate guarding by males, followed by facultative sharing of mating opportunities. While males in other species, such as chimpanzees, olive baboons, and lions, also collaborate to guard females against other males, the coalition-like behavior in stump-tailed macaques differs from them in the following aspects. In the case of lions, a group of females with a few males, called a pride, is formed and the males collectively defend the females from other invasive males. The alpha male mostly monopolizes the mating opportunities, and the subordinate allies have either no access to the females or only limited access not through active sharing (Bygott et al., 1979; Packer & Pusey, 1982). This is in contrast to the case of stump-tailed macaques, where a coalition-like unit is formed by a few males in a large multi-male multi-female group to exclude other males in the same group from mating competition, and the mating opportunities thus secured are actively shared within the unit. Similarly, in olive baboons, subordinate males form a coalition to jointly attack a dominant male, thereby increasing their future access to females. However, unlike stump-tailed macaques, olive baboons do not actively share copulations (Bercovitch, 1988). On the other hand, the observed case of a chimpanzee female copulating with eight males within a short time period (Watts, 1998) might be comparable to the facultative mate sharing in the stump-tailed macaque, although this was considered exceptional, observed only in the Ngogo population, which is considerably larger than the other populations (Watts, 1998).

A remarkable feature of the coalition-like behavior in stump-tailed macaques is that the alpha male appears to pay a reproductive cost by giving mating opportunities away to his allies, who in return offer collaborative work efforts, and as a result they gain reproductive advantage as a team. Another point deserving attention is that coalition formation manifested as joint aggression on a target may involve triadic awareness among the attacker, helper, and target (Harcourt & de Waal, 1992; Paxton et al., 2010; Schino, Tiddi, & Di Sorrentino, 2006; Silk, 1999), which is deemed to be cognitively more demanding than understanding the dyadic relationship between the self and another individual (Hemelrijk et al., 2013). For that matter, the coalition-like behavior in stump-tailed macaques, which sometimes involves collaboration among more than two individuals, may also require extra capacity of social cognition and that may be why similar behavior is rare in non-human animals.

Why do male stump-tailed macaques, unlike males of closely related species, exhibit this peculiar behavior? To put it in another way, what are the socio-ecological factors in stump-tailed macaques that may have favored the evolution of this behavior? Here, we propose that the absence of signs of ovulation in female stump-tailed macaques may be a key. In many primate species, females exhibit visual or olfactory signs of ovulation during the fertile period of the reproductive cycle. Conspicuous estrous signals such as sexual swellings enhance male-male competition, providing females more opportunities for mate choice (Nunn, 1999; Nunn, van Schaik, & Zinner, 2001; Zinner et al., 2004). Advertisement of female reproductive status is often seen in Old World monkeys living in multi-male multi-female societies, such as most macaques, baboons, and chimpanzees (Nunn, 1999; Nunn et al., 2001; Zinner et al., 2004). When female reproductive status is advertised, it is relatively easy for the alpha male to monopolize fertilizations, as in that case he can concentrate all his guarding efforts on the females fertile at that moment. On the other hand, when female ovulation is cryptic, the alpha male is no longer able to adopt the selective guarding strategy, and reproductive monopoly is only possible if all cycling females are guarded all the time. Our hypothesis is that the difficulty in establishing reproductive monopoly by the alpha male due to concealed ovulation may have promoted the formation of a coalition-like unit among dominant males.

Our discovery of the formation of a coalition-like unit, followed by active sharing of mating opportunities, in male stump-tailed macaques demands a revision of the existing socioecological models in primate social systems. As far as we are aware, this is the first documented case in non-human primates of collaborative effort for acquiring resources based on active sharing among the collaborators. We have hypothesized that the lack of estrous signs in female stump-tailed macaques, unlike many Old-World monkeys, is a key factor enhancing the coalition-like behavior. Concealed ovulation is likely to reduce the extent to which fertilizations are monopolized by dominant males. In our mathematical model, this effect is represented by the reduction in parameter *β*. The model predicts that male-male coalition is more likely to occur when *β* is small, confirming the logical consistency of our hypothesis. From the female’s perspective, monopolization by dominant males means limited opportunities for female mate choice, particularly when they prefer copulations with subordinate or out-group males. Thus, concealed ovulation may be considered as a female strategy to facilitate mate choice. Further extending the argument, the formation of coalition followed by active sharing of mating opportunities may be a counter strategy of dominant males. In other words, being unable to control female reproduction on his own, the alpha male may gain more by surrendering some fertilization opportunities to elicit cooperation by subordinates. Hence, the intensified sperm competition in stump-tailed macaques (García Granados et al., 2014) may be a joint consequence of female concealment of fertility states and male sharing of mating opportunities. In addition, a potentially relevant observation is that female stump-tailed macaques do not produce copulation calls (Blurton Jones & Trollope, 1968). Although the function of female copulation calls is still a matter of contention (Bernstein et al., 2016; Maestripieri & Roney, 2005), a possible interpretation is that female stump-tailed macaques do not make any effort to induce male mate guarding.

The present study has also revealed the importance of the number of males in a group as a predictor for the formation of coalitions among dominant males. In other words, male-male coalition is more likely to be formed when there are more males in a group. In our field site, we observed five groups of stump-tailed macaques consisting of 391 individuals, or on average 78.2 individuals per group. The relatively large group size is primarily due to the semi-provisioning conditions in our study site, and this factor also appears to affect the socioeconomic sex ratio, i.e., the ratio of the number of adult females to the number of adult males. The average socioeconomic sex ratio in our sample is 1.33, while those that have been previously reported for other populations of stump-tailed macaques are approximately 5.7 (Fooden, 1990). The smaller socioeconomic sex ratio indicates more intense male-male competition. Hence, both large number of males per group and small socioeconomic ratio may have facilitated the occurrence of coalition formation by dominant males in our study population.

While our model predicts monotonic increase of the coalition size with the increasing number of males in the group, we observed coalitions of two or three males, but never four or more. This discrepancy might indicate that there exist additional factors restricting the coalition size that are not considered in the model. A possible factor is, as mentioned earlier, the limited social cognition in non-human animals. Actually, psychological experiments on cooperative tasks have revealed that collaboration is possible among two or three chimpanzee subjects, but is much more difficult when four or more subjects are involved ((Hirata & Fuwa, 2007; Kaigaishi et al., 2019; Tomonaga et al., 2004)). For the recognition of quadradic relations, an individual has to recognize the possible combinations of dyadic and triadic relations, exponentially increasing the socio-cognitive loading in the brain.

We have also observed within-species variation in the extent to which copulations are monopolized by dominant males, which is represented by *β* in our model. Despite the marked ecological similarities between groups, the estimated *β*′ ranged from 0.30 to 0.97. In the Third (*β*′ = 0.97) and Wngklm (*β*′ = 0.78) groups, copulations were almost completely monopolized by the alpha males, a situation that is called “despotic.” This contrasts with the conventional classification of primate societies, in which stump-tailed macaques are characterized as having “egalitarian” societies (Matsumura, 1999), or class 3 social systems (Thierry et al., 2004). The traditional classification intends to place each species on a single position on the despotic-egalitarian spectrum, based largely on the species-level characterizations of ecological factors, such as whether or not a given species is seasonal breeder, or the abundance and spatial distribution of food resources (Sterck et al., 1997). However, our observations clearly suggest that the level of despotism as indicated by *β* is determined not necessarily in such a top-down manner, but in a more bottom-up way, such that it may vary within species according to the idiosyncrasies of each group. For example, our field observation indicates that the despotic nature of the Third group may have been caused not only by the physical strength of the alpha male, THR-M01, but by the absence of competent rivals; in fact, other males seem either too old or immature to challenge him. Therefore, it appears that bottom-up mechanisms determine *β* in each group, which then determines whether the alpha male will adopt the solo monopolization strategy or the coalition strategy.

Finally, our model predicts the future dynamics in the stump-tailed macaque groups. For example, when youngsters in the Third group become sufficiently mature to challenge the alpha male, and as a consequence *β* is reduced, our model predicts that the alpha male will form a coalition-like unit with other males. We expect that a longitudinal observation of wild stump-tailed macaques will confirm these model predictions. In conclusion, stump-tailed macaques are characterized by societies ranging from despotism to egalitarianism, and from monopolization of females by a dominant male to male-male coalition coupled with active sharing of mating opportunities. Future studies on wild stump-tailed macaques may shed new light on the origins and evolution of altruism and cooperation in mammalian societies, including the hyper-cooperation in human societies (Burkart et al., 2014).

## CONCLUSION

We have reported a novel observation of facultative sharing of mating opportunities among males in a wild population of stump-tailed macaques. Our observational data, mathematical analysis, and empirical test of model predictions altogether indicate that the observed behavior can be interpreted as a coalition-like behavior, in which dominant males collaboratively guard females from mating with subordinate males, and actively share the secured mating opportunities within the allies. The mathematical analysis predicts that less intense despotism and greater number of males in a group are to be associated with the coalition-like behavior. We have further argued that the lack of estrus signs in stump-tailed macaque females may be a key factor that accounts for the occurrence of the coalition-like behavior in this species.

## ACKNOWLEDGEMENTS

We thank Warayut Nilpaung and Chuchat Choklap, Phanlerd Inprasoet, Wanchai Inprasoet, Napatchaya Techaatiwatkun, Yuzuru Hamada, Ikuma Adachi, Takeshi Nishimura, Ikki Matsuda for their kind support. We got funding from JSPS KAKENHI (#16H04848, #16J0098), Young Science Explorer Grant of National Geographic Foundation for Science and Exploration – Asia, Kyoto University Foundation, the Cooperation Research Programs of Wildlife Research Center, the Sasakawa Scientific Research Grant from The Japan Science Society, Japan Science and Technology Agency Core Research for Evolutional Science and Technology 17941861 (#JPMJCR17A4), and Ministry of Education, Culture, Sports, Science and Technology, Grant-in-Aid for Scientific Research on Innovative Areas #4903 (Evolinguistic) 17H06380 and 17H06381.

## APPENSICS

We have Tables S1, S2, and Figure S1 in Supplementary Materials as separately attached.

## Notes

### Competing Interest Statement

The authors have declared no competing interest.

### Summary of Updates

changed title, main text and abstract revised

## REFERENCES

1. Altmann, S. A. (1962). A field study of the sociology of rhesus monkeys, Macaca mulatta. Annals of the New York Academy of Sciences, 102(2), 338–435. https://doi.org/10.1111/j.1749-6632.1962.tb13650.x

2. Bercovitch, F. B. (1988). Coalitions, cooperation and reproductive tactics among adult male baboons. Animal Behaviour, 36(4), 1198–1209. https://doi.org/10.1016/S0003-3472(88)80079-4

3. Bernstein, S. K., Sheeran, L. K., Wagner, R. S., Li, J.-H., & Koda, H. (2016). The vocal repertoire of Tibetan macaques (Macaca thibetana): A quantitative classification. American Journal of Primatology, 78(9), 937–949. https://doi.org/10.1002/ajp.22564

4. Bissonnette, A., de Vries, H., & van Schaik, C. P. (2009). Coalitions in male Barbary macaques, Macaca sylvanus: strength, success and rules of thumb. Animal Behaviour, 78(2), 329–335. https://doi.org/10.1016/j.anbehav.2009.05.010

5. Blurton Jones, N. G., & Trollope, J. (1968). Social behaviour of stump-tailed macaques in captivity. Primates, 9(4), 365–393.

6. Brereton, A. R. (1994). Copulatory behavior in a free-ranging population of stumptail macaques (Macaca arctoides) in Mexico. Primates, 35(2), 113–122. https://doi.org/10.1007/BF02382048

7. Burkart, J. M., Allon, O., Amici, F., Fichtel, C., Finkenwirth, C., Heschl, A., Huber, J., Isler, K., Kosonen, Z. K., Martins, E., Meulman, E. J., Richiger, R., Rueth, K., Spillmann, B., Wiesendanger, S., & van Schaik, C. P. (2014). The evolutionary origin of human hyper-cooperation. Nature Communications, 5(1), 4747. https://doi.org/10.1038/ncomms5747

8. Bygott, J. D., Bertram, B. C. R., & Hanby, J. P. (1979). Male lions in large coalitions gain reproductive advantages. Nature, 282(5741), 839–841. https://doi.org/10.1038/282839a0

9. Cerda-Molina, A. L., Hernández-López, L., Rojas-Maya, S., Murcia-Mejía, C., & Mondragón-Ceballos, R. (2006). Male-induced sociosexual behavior by vaginal secretions in Macaca arctoides. International Journal of Primatology, 27(3), 791–807. https://doi.org/10.1007/s10764-006-9045-0

10. Chapais, B. (1995). Alliances as a means of competition in primates. American Journal of Physical Anthropology, 38, 115–136.

11. Estep, D. Q., Bruce, K. E. M., Johnston, M. E., & Gordon, T. P. (1984). Sexual Behavior of Group-Housed Stumptail Macaques (Macaca arctoides): Temporal, Demographic and Sociosexual Relationships. Folia Primatologica, 42(2), 115–126. https://doi.org/10.1159/000156154

12. Fooden, J. (1990). The bear macaque, Macaca arctoides: a systematic review. Journal of Human Evolution, 19(6–7), 607–686. https://doi.org/10.1016/0047-2484(90)90002-S

13. García Granados, M. D., Hernández López, L. E., Córdoba Aguilar, A., Cerda Molina, A. L., Pérez-Ramírez, O., & Mondragón-Ceballos, R. (2014). Effect of photoperiod on characteristics of semen obtained by electroejaculation in stump-tailed macaques (Macaca arctoides). Primates, 55(3), 393–401. https://doi.org/10.1007/s10329-014-0414-6

14. Harcourt, A. H., de Waal, F. B. M., & others. (1992). Coalitions and alliances in humans and other animals. Oxford University Press Oxford.

15. Hemelrijk, C. K., Puga-Gonzalez, I., & Steinhauser, J. (2013). Cooperation, Coalition, and Alliances. In Handbook of Paleoanthropology (Issue Hamilton 1964, pp. 1–27). Springer Berlin Heidelberg. https://doi.org/10.1007/978-3-642-27800-6_43-4

16. Hirata, S., & Fuwa, K. (2007). Chimpanzees (Pan troglodytes) learn to act with other individuals in a cooperative task. Primates, 48(1), 13–21. https://doi.org/10.1007/s10329-006-0022-1

17. Kaigaishi, Y., Nakamichi, M., & Yamada, K. (2019). High but not low tolerance populations of Japanese macaques solve a novel cooperative task. Primates, 60(5), 421–430. https://doi.org/10.1007/s10329-019-00742-z

18. Kutsukake, N., & Nunn, C. L. (2006). Comparative tests of reproductive skew in male primates: the roles of demographic factors and incomplete control. Behavioral Ecology and Sociobiology, 60(5), 695–706. https://doi.org/10.1007/s00265-006-0213-1

19. Maestripieri, D., & Roney, J. R. (2005). Primate copulation calls and postcopulatory female choice. Behavioral Ecology, 16(1), 106–113. https://doi.org/10.1093/beheco/arh120

20. Matsumura, S. (1999). The evolution of “egalitarian” and “despotic” social systems among macaques. Primates, 40(1), 23–31. https://doi.org/10.1007/BF02557699

21. Noë, R., & Sluijter, A. A. (1995). Which adult male savanna baboons form coalitions? International Journal of Primatology, 16(1), 77–105. https://doi.org/10.1007/BF02700154

22. Nunn, C. L, van Schaik, C. P., & Zinner, D. (2001). Do exaggerated sexual swellings function in female mating competition in primates? A comparative test of the reliable indicator hypothesis. Behavioral Ecology, 12(5), 646–654. https://doi.org/Article

23. Nunn, Charles L. (1999). The evolution of exaggerated sexual swellings in primates and the graded-signal hypothesis. Animal Behaviour, 58(2), 229–246. https://doi.org/10.1006/anbe.1999.1159

24. Packer, C., & Pusey, A. E. (1982). Cooperation and competition within coalitions of male lions: kin selection or game theory? Nature, 296(5859), 740–742. https://doi.org/10.1038/296740a0

25. Pandit, S. A., Pradhan, G. R., & van Schaik, C. P. (2020). Why Class Formation Occurs in Humans but Not among Other Primates: A Primate Coalitions Model. Human Nature, 31(2), 155–173. https://doi.org/10.1007/s12110-020-09370-9

26. Pandit, S. A., & van Schaik, C. P. (2003). A model for leveling coalitions among primate males: toward a theory of egalitarianism. Behavioral Ecology and Sociobiology, 55(2), 161–168. https://doi.org/10.1007/s00265-003-0692-2

27. Paxton, R., Basile, B. M., Adachi, I., Suzuki, W. A., Wilson, M. E., & Hampton, R. R. (2010). Rhesus monkeys (Macaca mulatta) rapidly learn to select dominant individuals in videos of artificial social interactions between unfamiliar conspecifics. Journal of Comparative Psychology, 124(4), 395–401. https://doi.org/10.1037/a0019751

28. Schino, G., Tiddi, B., & Di Sorrentino, E. P. (2006). Simultaneous classification by rank and kinship in Japanese macaques. Animal Behaviour, 71(5), 1069–1074. https://doi.org/10.1016/j.anbehav.2005.07.019

29. Silk, J. B. (1999). Male bonnet macaques use information about third-party rank relationships to recruit allies. Animal Behaviour, 58(1), 45–51. https://doi.org/10.1006/anbe.1999.1129

30. Smith, J. E., Van Horn, R. C., Powning, K. S., Cole, A. R., Graham, K. E., Memenis, S. K., & Holekamp, K. E. (2010). Evolutionary forces favoring intragroup coalitions among spotted hyenas and other animals. Behavioral Ecology, 21(2), 284–303. https://doi.org/10.1093/beheco/arp181

31. Sterck, E. H. M., Watts, D. P., & van Schaik, C. P. (1997). The evolution of female social relationships in nonhuman primates. Behavioral Ecology and Sociobiology, 41(5), 291–309. https://doi.org/10.1007/s002650050390

32. Thierry, B., Singh, M., & Kaumanns, W. (2004). Macaque societies: a model for the study of social organization (Vol. 41). Cambridge University Press.

33. Tomonaga, M., Tanaka, M., Matsuzawa, T., Myowa-Yamakoshi, M., Kosugi, D., Mizuno, Y., Okamoto, S., Yamaguchi, M. K., & Bard, K. A. (2004). Development of social cognition in infant chimpanzees (Pan troglodytes): Face recognition, smiling, gaze, and the lack of triadic interactions. Japanese Psychological Research, 46(3), 227–235. https://doi.org/10.1111/j.1468-5584.2004.00254.x

34. Toyoda, A., & Malaivijitnond, S. (2018). The First Record of Dizygotic Twins in Semi-Wild Stump-Tailed Macaques (Macaca arctoides) Tested Using Microsatellite Markers and the Occurrence of Supernumerary Nipples. Mammal Study, 43(3), 207–212. https://doi.org/10.3106/ms2017-0081

35. Toyoda, A., Maruhashi, T., Malaivijitnond, S., & Koda, H. (2017). Speech-like orofacial oscillations in stump-tailed macaque (Macaca arctoides) facial and vocal signals. American Journal of Physical Anthropology, 164(2), 435–439. https://doi.org/10.1002/ajpa.23276

36. van Schaik, C. P., Pandit, S. A., & Vogel, E. R. (2004). A model for within-group coalitionary aggression among males. Behavioral Ecology and Sociobiology, 57(2), 101–109. https://doi.org/10.1007/s00265-004-0818-1

37. van Schaik, C. P., Pandit, S. A., & Vogel, E. R. (2006). Toward a general model for male-male coalitions in primate groups. In Cooperation in Primates and Humans: Mechanisms and Evolution (pp. 151–171). Springer Berlin Heidelberg. https://doi.org/10.1007/3-540-28277-7_9

38. Van Schaik, C. P., Pandit, S. A., & Vogel, E. R. (2006). Toward a general model for male-male coalitions in primate groups. In Cooperation in Primates and Humans: Mechanisms and Evolution (pp. 151–171). https://doi.org/10.1007/3-540-28277-7_9

39. Watts, D. P. (1998). Coalitionary mate guarding by male chimpanzees at Ngogo, Kibale National Park, Uganda. Behavioral Ecology and Sociobiology, 44(1), 43–55. https://doi.org/10.1007/s002650050513

40. Widdig, A., Streich, W. J., & Tembrock, G. (2000). Coalition formation among male Barbary macaques (Macaca sylvanus). American Journal of Primatology, 50(1), 37–51. https://doi.org/10.1002/(SICI)1098-2345(200001)50:1<37::AID-AJP4>3.0.CO;2-3

41. Young, C., Schülke, O., & Ostner, J. (2014). How males form coalitions against group rivals and the Pandit/van Schaik coalition model. Behaviour, 151(7), 907–934. https://doi.org/10.1163/1568539X-00003166

42. Zinner, D., van Schaik, C. P., Nunn, C. L., & Kappeler, P. M. (2004). Sexual selection and exaggerated sexual swellings of female primates. In Sexual Selection in Primates: New and Comparative Perspectives (pp. 71–89). https://doi.org/10.1002/evan.10023

