## Supplementary material for "Mate sharing in male stump-tailed macaques as a possible case of coalition-like behavior to modify the group-wise fitness distribution": Tables S1 to S2 and Figure S1 (from another version of this preprint: version 3, 2021-02-08)

Supplementary Materials for

**Sharing mating opportunities by coalition males in stump-tailed macaques: elucidation of ecological mechanism by diversion of PvS model**

Aru Toyoda*, Tamaki Maruhashi, Suchinda Malaivijitnond, Yasuo Ihara*, Hiroki Koda*.

**This PDF file includes:**

Tables S1 to S2

Figure S1

Table S1.

| Group | Adult males | Adult females | Infant^1^ | Unidentified^2^ | Total |
| --- | --- | --- | --- | --- | --- |
| Ting | 30 | 33 | 39 | 18 | 120 |
| Nadam | 20 | 30 | 21 | 4 | 75 |
| Third | 23 | 23 | 20 | 13 | 79 |
| Fourth | 16 | 27 | 22 | 9 | 74 |
| Wngklm | 8 | 11 | 12 | 12 | 43 |
| Total | 97 | 124 | 114 | 56 | 391 |

Age-sex composition of each group in Khao Krapuk Khao Taomor non-hunting area.

^1^Infants were 0−2 year-old monkeys. ^2^Unidentified monkeys were all subadult monkeys who were not stably observed in the group, the numbers of which have been reported above.

Table S2.

**Table S2-1.** All serial copulation bouts of Ting group

| **Bout-id** | **Group** | **Observed date** | **Female** | **Male: TNG-M02** | **Male: TNG-M07** | **Male: TNG-M04** | **Male: TNG-M08** | **Male: TNG-M09** | **Male: TNG-M30** | **Male: TNG-M18** | **Male: TNG-M03** | **Male: TNG-M17** | **Total (N)** |
| --- | --- | --- | --- | --- | --- | --- | --- | --- | --- | --- | --- | --- | --- |
| **ting-1** | Ting | 2015/10/23 | TNG-F11 | 7(4) | 4(4) | - | - | - | - | - | 1(0) | - | 12(8) |
| **ting-2** | Ting | 2015/11/8 | TNG-F23 | 8(8) | 6(6) | - | - | - | - | - | - | - | 14(14) |
| **ting-3** | Ting | 2015/12/30 | TNG-F05 | 1(1) | 3(3) | - | - | 2(2) | - | - | - | - | 6(6) |
| **ting-4** | Ting | 2016/1/28 | TNG-F10 | - | - | - | - | 1(0) | - | 2(2) | - | 1(0) | 4(2) |
| **ting-5** | Ting | 2016/10/13 | TNG-F18 | 6(6) | 1(1) | - | - | - | - | - | - | - | 7(7) |
| **ting-6** | Ting | 2016/10/13 | TNG-F18 | 4(4) | - | - | - | - | - | - | - | - | 4(4) |
| **ting-7** | Ting | 2016/11/3 | TNG-F32 | - | - | 4(3) | 2(0) | - | 2(1) | - | - | - | 8(4) |
| **ting-8** | Ting | 2017/2/1 | TNG-F09 | 3(3) | 1(0) | - | - | - | - | - | - | - | 4(3) |
| **ting-9** | Ting | 2017/2/8 | TNG-F11 | 0 | 4(3) | - | 1(1) | - | - | - | - | - | 5(4) |

**Table S2-2.** All serial copulation bouts of Nadam group

| **Bout-id** | **Group** | **Observed date** | **Female** | **Male: NDM-M01** | **Male: NDM-M03** | **Total** |
| --- | --- | --- | --- | --- | --- | --- |
| **nadam-1** | Nadam | 2015/12/19 | NDM-F07 | 4(4) | - | 4(4) |
| **nadam-2** | Nadam | 2016/2/9 | NDM-F12 | 3(3) | 3(3) | 6(6) |
| **nadam-3** | Nadam | 2016/6/14 | NDM-F01 | 3(3) | 2(2) | 5(5) |
| **nadam-4** | Nadam | 2016/6/23 | NDM-F25 | 4(4) | 2(1) | 6(5) |

**Table S2-3.** All serial copulation bouts of Third group

| **Bout-id** | **Group** | **Observed date** | **Female** | **Male: TRD-M01** | **Total** |
| --- | --- | --- | --- | --- | --- |
| **third-1** | Third | 2015/12/24 | TRD-F25 | 16(16) | 16(16) |
| **third-2** | Third | 2016/10/3 | TRD-F24 | 31(31) | 31(31) |
| **third-3** | Third | 2016/10/12 | TRD-F16 | 4(1) | 4(1) |
| **third-4** | Third | 2016/11/14 | TRD-F16 | 6(1) | 6(1) |
| **third-5** | Third | 2017/1/18 | TRD-F29 | 8(8) | 8(8) |

**Table S2-4.** All serial copulation bouts of Fourth group

| **Bout-id** | **Group** | **Observed date** | **Female** | **Male: FTH-M01** | **Male: FTH-M02** | **Male: FTH-M15** | **Total** |
| --- | --- | --- | --- | --- | --- | --- | --- |
| **fourth-1** | Fourth | 2015/10/10 | FTH-F13 | 8(8) | 1(1) | - | 9(9) |
| **fourth-2** | Fourth | 2015/10/24 | FTH-F13 | 11(11) | 3(3) | - | 14(14) |
| **fourth-3** | Fourth | 2016/6/16 | FTH-F08 | - | 7(6) | - | 7(6) |
| **fourth-4** | Fourth | 2016/7/21 | FTH-F02 | 4(4) | - | - | 4(4) |
| **fourth-5** | Fourth | 2016/7/23 | FTH-F11 | 4(4) | - | - | 4(4) |
| **fourth-6** | Fourth | 2016/10/25 | FTH-F11 | 2(2) | 1(1) | 7(7) | 10(10) |

**Table S2-5.** All serial copulation bouts of Wngklm group

| **Bout-id** | **Group** | **Observed date** | **Female** | **Male: WKM-M01** | **Total** |
| --- | --- | --- | --- | --- | --- |
| **wngklm-1** | Wngklm | 2016/2/2 | WKM-F04 | 6(6) | 6(6) |
| **wngklm-2** | Wngklm | 2016/8/23 | WKM-F11 | 9(9) | 9(9) |

All rows in the tables represent a single copulation bout and indicate the “Group”, “Observed Dates”, “Female ID”, and “Male ID” involved in the bout. The numbers represent the count numbers of copulation event (in parentheses are the number of copulations with ejaculation).

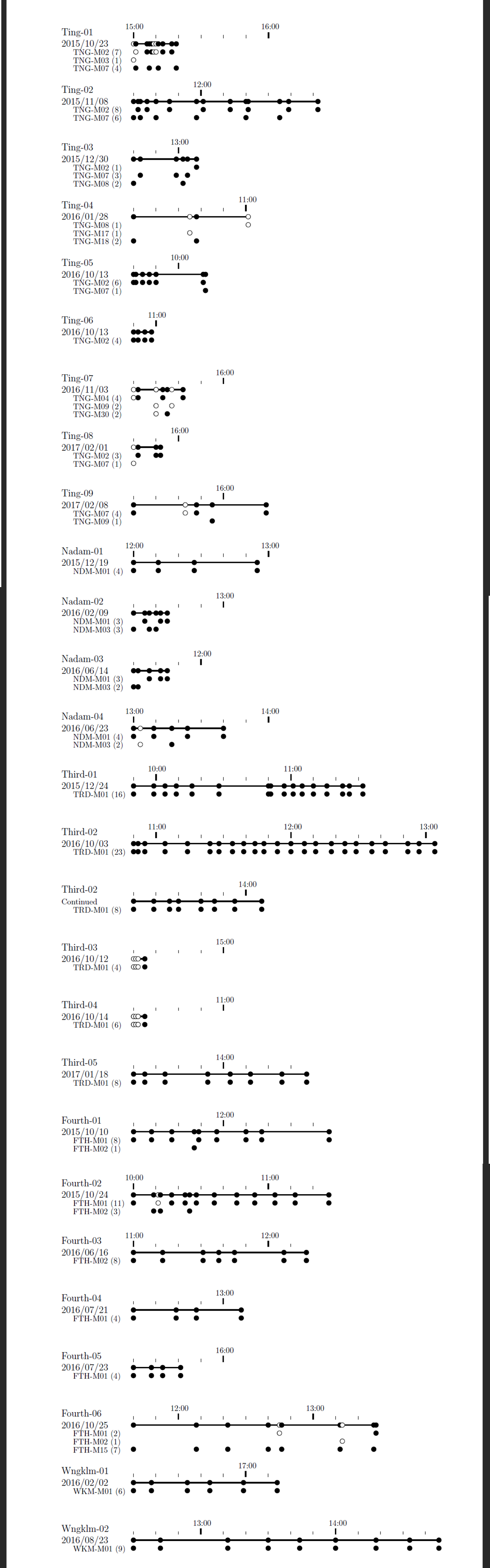

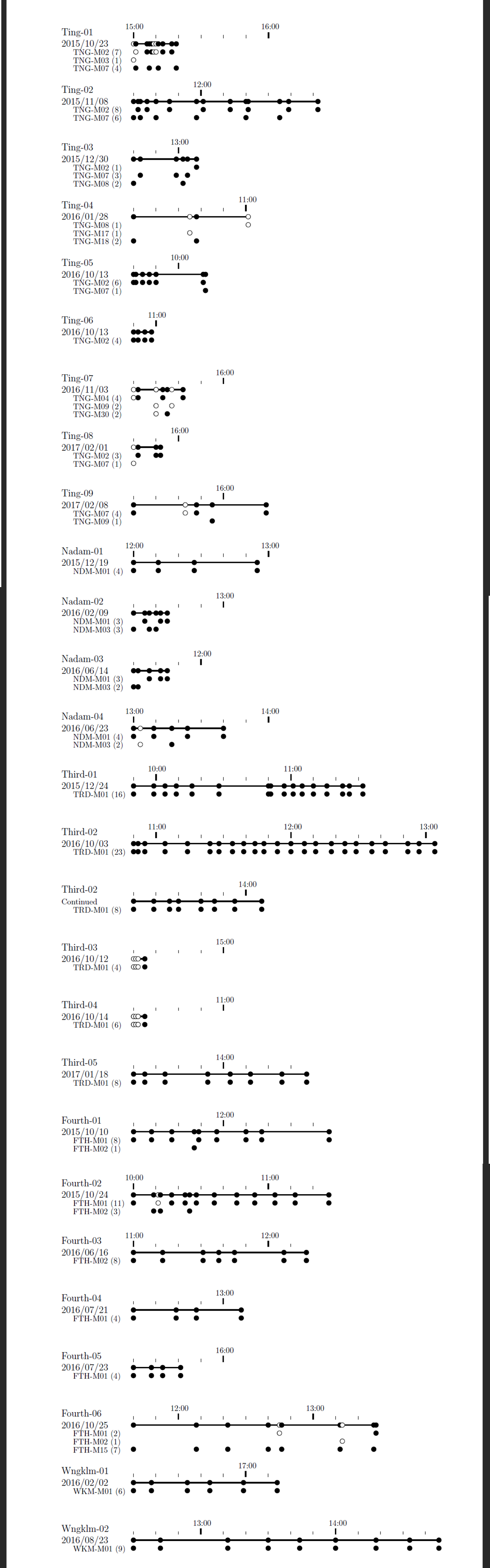

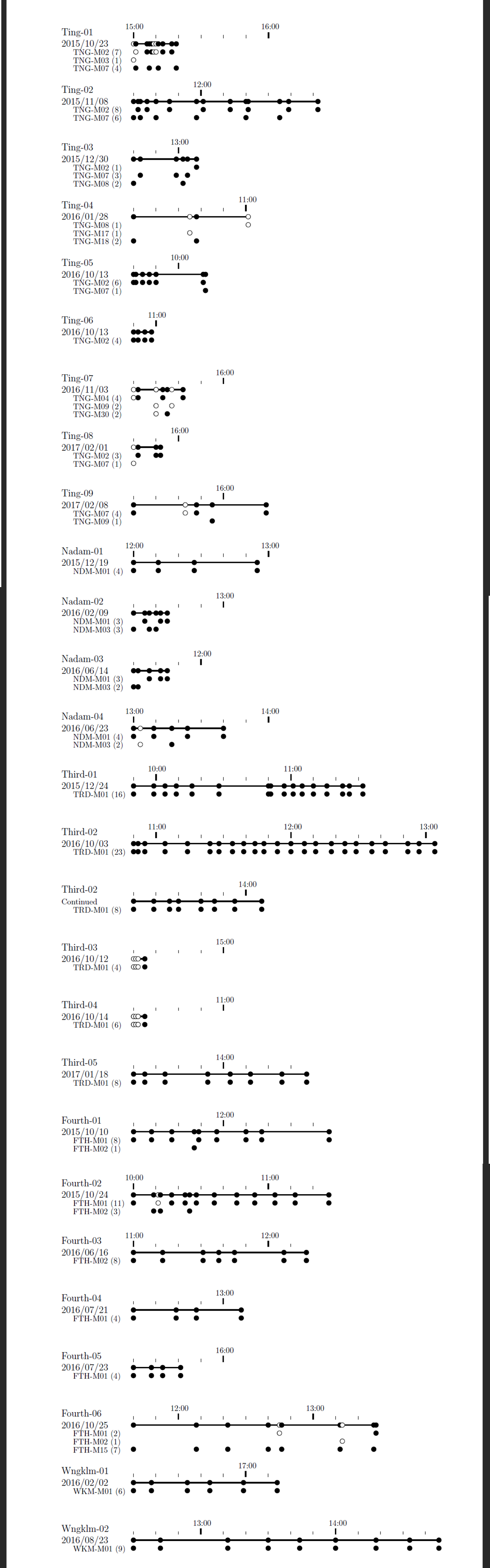

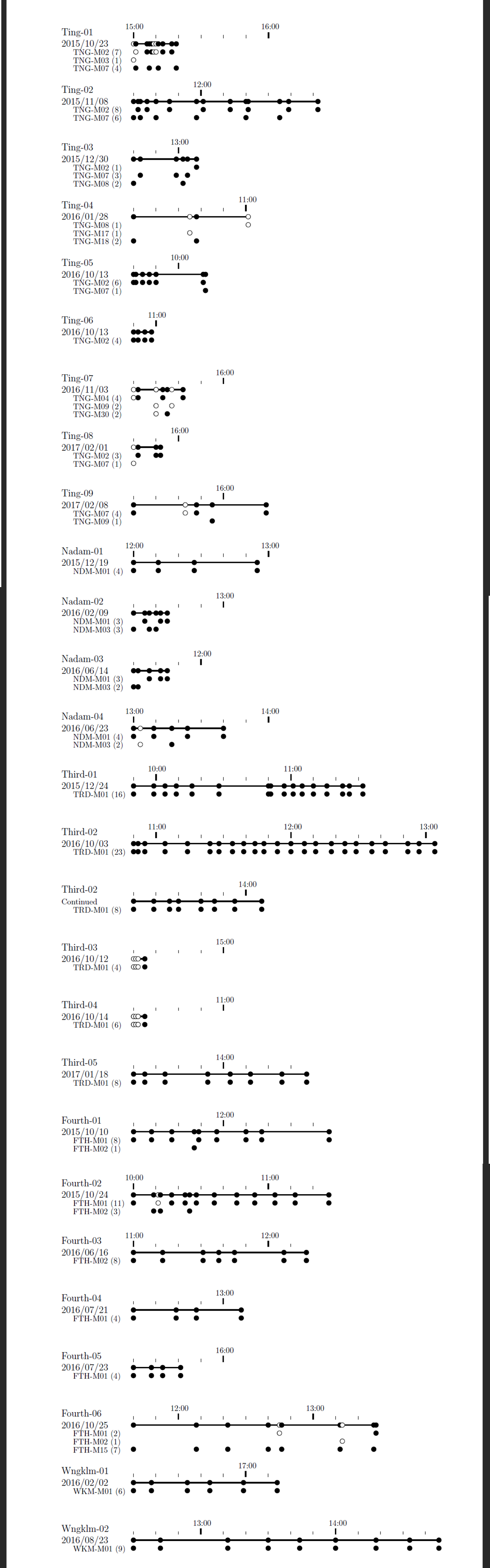

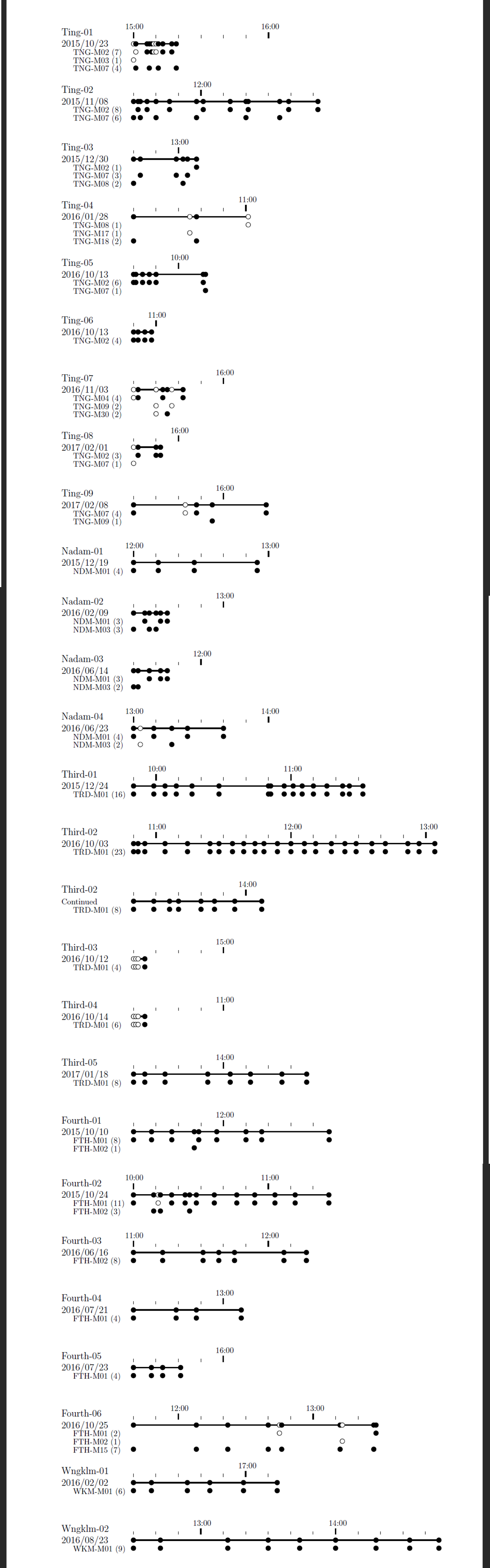

**Figure S1**

Sequential figure for each event of Serial copulations. Black circles indicate copulations end with ejaculation, and white circles indicate copulations end without ejaculation.
